# Adaptive and Wireless Recordings of Electrophysiological Signals during Concurrent Magnetic Resonance Imaging

**DOI:** 10.1101/259762

**Authors:** Ranajay Mandal, Nishant Babaria, Jiayue Cao, Zhongming Liu, Senior Member IEEE

**Affiliations:** Weldon School of Biomedical Engineering and Purdue Institute for Integrative Neuroscience, Purdue University, West Lafayette, IN, USA; School of Electrical and Computer Engineering, Purdue University, West Lafayette, IN, USA

**Keywords:** Simultaneous fMRI-EEG, Wireless, frequency modulation, gradient artifacts

## Abstract

Strong electromagnetic fields that occur during functional magnetic resonance imaging (fMRI) presents a challenging environment for concurrent electrophysiological recordings. Here, we present a miniaturized, wireless platform – “MR-Link” (Multimodal Recording Link) that provides a hardware solution for simultaneous electrophysiological and fMRI signal acquisition. The device detects the changes in the electromagnetic field during fMRI to synchronize amplification and sampling of electrophysiological signals with minimal artifacts. It wirelessly transmits the recorded data at a frequency detectable by the MR-receiver coil. The transmitted data is readily separable from MRI in the frequency domain. To demonstrate its efficacy, we used this device to record electrocardiograms and somatosensory evoked potential during concurrent fMRI scans. The device minimized the fMRI-induced artifacts in electrophysiological data and wirelessly transmitted the data back to the receiver coil without compromising fMRI signal quality. The device is compact (22 mm dia., 2gms) and can be placed within the MR-bore to precisely synchronize with fMRI. Therefore, MR-Link offers an inexpensive system by eliminating the need for amplifiers with a high dynamic range, high-speed sampling, additional storage or synchronization hardware for electrophysiological signal acquisition. It is expected to enable a broader range of applications of simultaneous fMRI and electrophysiology in animals and humans.

## I. Introduction

Simultaneous acquisition of functional magnetic resonance imaging (fMRI) in combination with electroencephalography (EEG), electrocorticography (ECoG), local field potentials (LFP), and single or multi-unit activity (SUA/MUA) holds great potential to bridge brain activity across spatial and temporal scales [1]–[10]. Despite its scientific premise and clinical potential, concurrent electrophysiological (EP) and MRI acquisition is challenging, since the MRI apparatus presents a hostile environment for recording bioelectric signals [3], [5], [8], [11]. MR-safety and compatibility are of paramount concern for any device working inside the MRI [12]–[14]. Apart from safety, the major bottleneck is the electromagnetic (EM) artifacts generated by (a) the static magnetic field, (b) strong RF deposition, and most importantly, (c) the rapidly changing gradient magnetic fields [6], [15]–[17]. These artifacts may be orders of magnitude larger than the signals of interest [15], [18]–[20]. As a result, the recorded EP data is often unusable to researchers without specialized technical expertise.

Widely used methods to remove EM artifacts involves sophisticated, offline post-processing of the artifact-corrupted dataset [3], [19]–[22]. Since these artifacts are often considerably larger than the signals of interest, existing systems all use highpower amplifiers with a large dynamic range and a high sampling rate (>20KHz) to accurately sample the corrupted signal [21], [23], [24]. Such amplifiers are frequently used along with additional shielding and powering modules, making the whole system complex, expensive, and of safety concern, especially in high-field MRI [17], [24]–[27]. Moreover, retrospective artifact removal is technically demanding, even if the artifacts and signals are recorded with high fidelity [28]–[31]. The artifacts may be non-stationary or inter-mixed with other types of noise (e.g. head motion) and thus difficult to fully isolate or remove [11], [20], [21], [28]–[32]. Even a small percentage of residual artifacts may be very problematic for recovering weak signals such as EEG [29].

Although rare, hardware solutions have been proposed to prevent EM artifacts from corrupting EP recordings. For example, Anami et al. have proposed a so-called “stepping stone sampling” strategy to skip the EM artifact by selectively sampling EEG only during the “silent” period of MRI [15]. This strategy requires the precise synchronization of the MR-scanner operational clocks and the EEG recording system, preexisting knowledge of the MRI pulse sequence, and a high fidelity recording system, all of which add a considerable level of complexity and thus limit the scope of application [15]. In addition, Hanson et al. have proposed a complementary strategy to wirelessly transmit EP recordings for reception by the MRI receiver coil, while utilizing the surplus hardware in the MRI system [33]. While this strategy reduced gradient artifacts, additional post-processing effort was still needed, e.g. to remove cross talks between “MR” and “Non-MR” data. Furthermore, they used a relatively bulky system (2 kg), which could only be operated from outside the MR-bore and require long wired connections, thereby rendering the signal acquisition susceptible to movement related artifacts.

These circumstances call for a much simpler and more robust method for EP recording alongside conventional fMRI. To meet this need, we developed a wireless and miniaturized device, namely “MR-Link” (Multimodal Recording Link), for high-fidelity EP recording during simultaneous fMRI. The device contained onboard coils, analog and digital circuits to detect the timing of gradient changes. Utilizing this timing information, the targeted EP signal was amplified and sampled discretely to avoid the EM artifacts. The device contained an onboard transmitter to send the recorded EP signals to the MRI receiver coil along with fMRI images (Figure 1). Thus, the device could operate locally and wirelessly, increasing the signal to noise ratio (SNR) while reducing the EM and movement artifacts [17], [34]. As an accessory to MRI, MR-Link facilitates the use of MRI scanner for both imaging and recording through its existing hardware. Hereafter, we describe the system design and implementation and a series of experimental results, which demonstrate the efficacy and potential of this device.

**Fig. 1.**
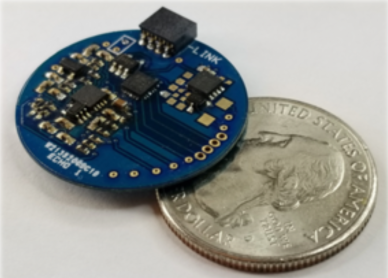
MR-Link device with a small footprint (22 mm diamter; 2 grams).

## II. Materials and Methods

### A. Device Operation

The device was designed to utilize the existing MR-hardware and the EM environment to trigger EP recording and wirelessly transmit digitized data. The device recorded the EP signal based on the input it received from the gradient detection system which monitored changes in the magnetic field inside the MRI bore [35]–[37]. The device amplified and filtered the EP signal and transmitted it wirelessly to the MR-receiver coil during MRI acquisition. The EP signal was modulated at a different frequency to ensure MR-signal and non-MR-signals were separated and demodulated without mutual interference (Figure 2). Figure 3 illustrates the major components of the device. Each of these components and their working principle are described in the following subsections.

**Fig. 2.**
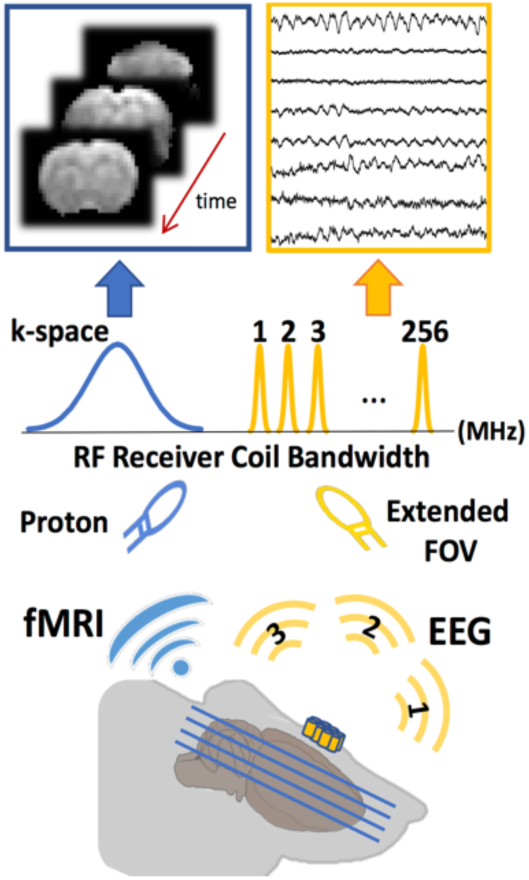
Concurrent fMRI and EP signal recording utilizing MR-receive coil. Amplified and digitized EP data is modulated by the MR-link recorder at discrete frequencies, visible to the MR-receiver coil. The MR signal from the subject is sensed by the receiver coil near the center frequency of MRI scanner (Blue). Simultaneously, the EP data, modulated with an offset (w.r.t the MRI center frequency) is also detected (Yellow). This spectrally isolated EP data becomes embedded into the extended FOV of the reconstructed MR image.

**Fig. 3.**
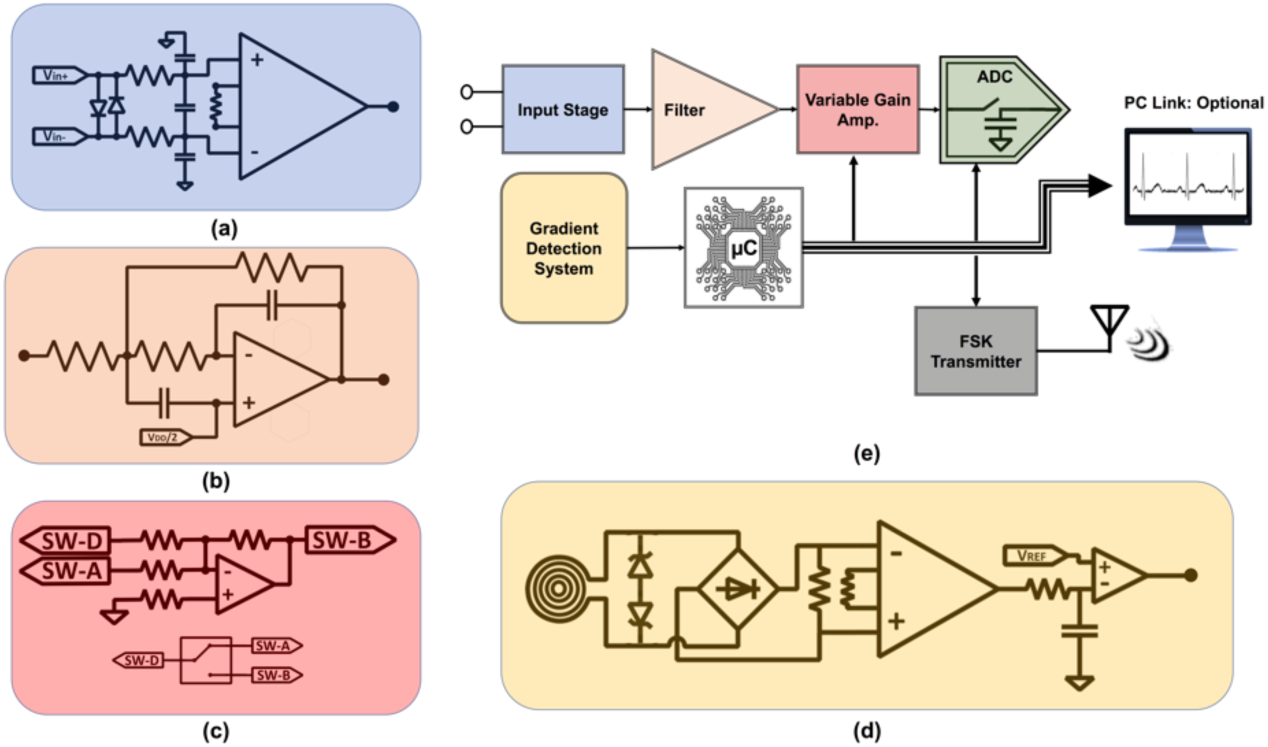
Simplified circuit schematic of MR-Link. The EP signal is processed by the analog filtering circuit (a,b), variable-gain amplifier (c), and subsequently digitized through the microcontroller (e). Gradient detection circuit (d) provides trigger information for variable-gain amplification and digitization. Digitized data is transmitted wirelessly through a low-power ultra high frequency (UHF) transmitter. Alternatively, the data can also be accessed through a USB-UART (Universal Asynchronous Receiver-Transmitter) PC-link. This PC-Link also provides the necessary power (<15mw required for analog processing and digitization). (a) Input differential stage with a low-pass filter (15 KHz) and gain of 6.02 dB. (b) Second order low-pass filter (15 KHz) with a gain of 0 dB. (c) Gain switching stage with gain of 54 dB during ‘plateau period’ and attenuation of −54 dB during ‘ramping period’ of the gradient. (d) Gradient detection circuit which monitors the electromagnetic field changes inside the MRI bore through a machine wound coil (14mm dia.) and outputs a binary signal to denoting the presence of gradient and RF artifacts.

#### 1.) Gradient detection circuit

The gradient detection circuit utilized a machine-wound copper coil to pick up variations in the magnetic field, as trapezoidal gradients were played out during fMRI. The differential signal from the coil was passed through a rectifier circuit and converted to a single-ended signal by an instrumentation amplifier. The circuit input further incorporated a voltage limiter circuit to limit the signal to ±3.6V. The single-ended signal was further filtered through a single-stage low-pass filter (<30kHz) (Figure 3(d)). The filter removed high frequency signal components to help avoid false triggers. A comparator was ultimately used to generate a negative logic binary output (hereafter referred to as the gradient trigger), reporting whether the magnetic field was static (binary ‘1’) or varying (binary ‘0’) (see Figure 4, 2^nd^ row). Thus, the gradient trigger signal was a ‘0’ during the ramping periods of a trapezoidal gradient and a ‘1’ during the plateau period. The gradient trigger was then fed into a microcontroller (µC) which managed the onboard operations of the device discussed hereafter.

**Fig. 4.**
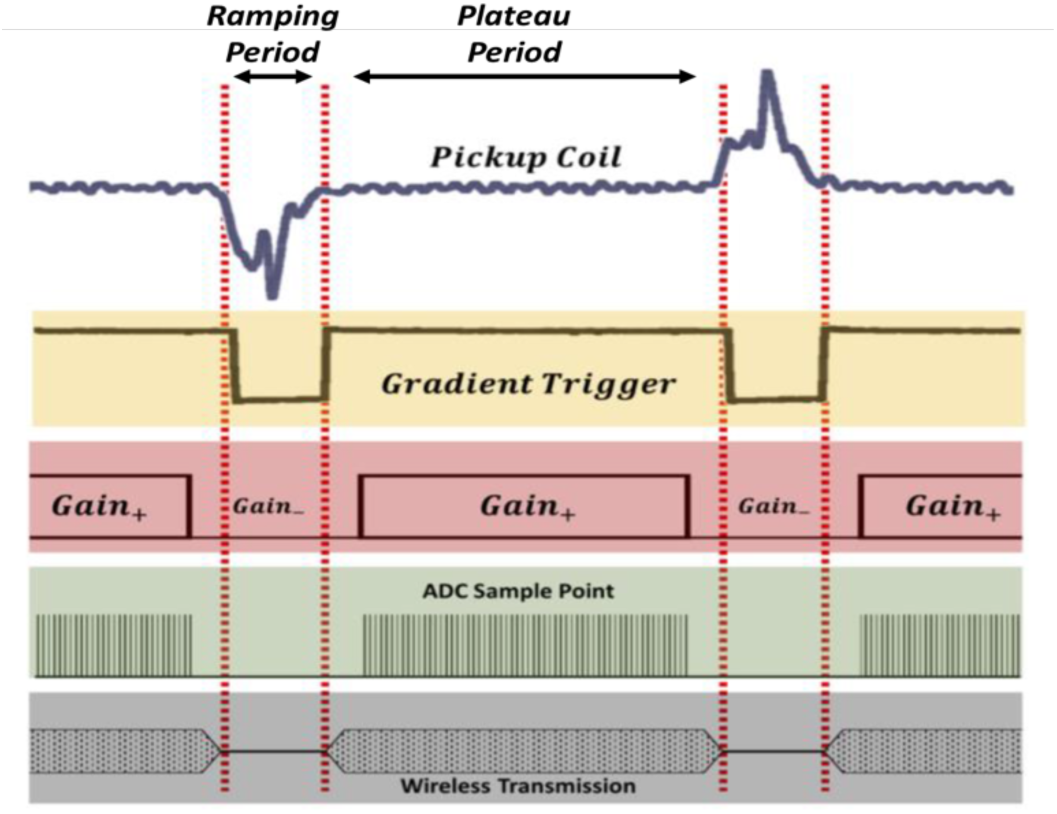
Gain switching, adaptive sampling, and data transmission driven by the gradient detection circuit. Pickup coil signal is translated into the binary gradient output which allows the onboard microcontroller to modulate gain and sample EP signal to avoid MR artifacts (Row 1-4). Digitized data is transmitted (wirelessly or wired) during ‘plateau period’ (Row 5).

#### 2.) Adaptive Sampling

Through an onboard µC, the gradient detection circuit made it possible to avoid gradient artifacts by sampling EP signals at discrete periods [15], [36], [37]. The µC learned the durations of the gradient plateau and ramping periods from the gradient triggers. The µC timestamped each edge of the gradient trigger, and calculated the duration of the plateau or ramping period. Based on this timing information, the µC switched the gain of analog amplification and applied analog-to-digital conversion (ADC). As a result, the recorded signal was amplified and digitized only when the gradient trigger was high, i.e. during the plateau period of the gradient, whereas it was attenuated during the ramping periods to prevent the gradient artifacts from saturating the amplifier.

The µC learned the regularity of gradients and predicted the occurrence of gradient ramping and plateau ahead of actual detection. The predictive control by the µC allowed the analog circuit to attenuate the artifact before it occurred. The µC continuously updated its prediction based on the most recent history about the timing of the detected gradients. The µC also scheduled other operations such as digitization and transmission with a 1MHz operational clock.

#### 3.) Analog filtering and amplification

The analog filtering and amplification system (or the analog circuit) on the device consisted of two variable gain amplifiers (VGA), a passive differential low-pass filter and another second-order active low-pass filter. Additionally, a voltage limiter circuit was incorporated at the input stage to limit the input signal to ±0.3V (Figure 3(a)). The main operation of the analog circuit was to attenuate the incoming signal when the gradient trigger was low and to provide amplification when the gradient trigger was high. This assured that the analog system did not saturate due to the strong gradient artifacts.

The low-pass filters were both cutoff at 15kHz to reject any high-frequency EM interference, while providing a sufficient bandwidth to capture a wide range of EP signals. The discrete-time variable-gain amplification was essential to subdue any gradient artifacts and prevent amplifier saturation. The two VGAs individually provided +/27 dB based on the gradient trigger. The maximum gain applied to the EP signal was 54 dB during the plateau periods and conversely, the maximum attenuation applied was −54 dB during the ramping periods. Apart from the VGAs, 6 dB of constant gain was provided by the instrumentation amplifier at the input stage, where the differential EP signal was converted to a single-ended signal for filtering and further processing by the VGAs (Figure 3(c)).

#### 4.) Wireless Transmission

The digitized packets were wirelessly transmitted through a low power ultra-high frequency (UHF) transmitter using binary Frequency Shift Keying (FSK). The carrier frequency of the RF signal was programmed within the bandwidth of the MR receiver coil, but outside of the frequency range for MR signal reception (Figure 2) [33], [36], [37]. The transmission power, carrier frequency (300.043MHz), deviation frequency (9.6KHz), data modulation types (FSK) and data baud rate (19.2Kbps) were all programmed and controlled through the µC. The built-in fractional-N PLL (phase-locked loop) of the transmitter was utilized for narrow-band operation. The reference frequency was generated through a crystal oscillator with high frequency (10ppm) and temperature stability (10ppm/°C^2^). Furthermore, any frequency error in crystal reference frequency were automatically corrected for through an inbuilt compensation register, which could also be adjusted manually to generate any specific output frequency with <1 ppm error. Additionally, the output power of the transmitter could be programmed from −16dBm to +14dBm with 0.4dBm resolution. This provided additional flexibility for the transmitted data to fit the RF amplifier’s dynamic range and be suitable for various receiver gain settings. Lastly, a fifth order Chebyshev filter was applied to the transmitter output to attenuate the 3rd and 5th order harmonics.

#### 5.) FOV extension for EP reception

The FOV for imaging was extended along the readout direction to allow RF-modulated EP data to be received by the MR-receiver coil. The image matrix along the readout direction contained 256 points. The receiver bandwidth was set to 333.333 KHz to provide a sufficient sampling rate for implementing FSK modulation of transmitted data. During each k-space line acquisition, the receive coil was ‘ON’ (set to acquire RF signals). The digitized EP data was transmitted during this time to embed itself in the extended FOV of the MR image (Figure 5).

**Fig. 5.**
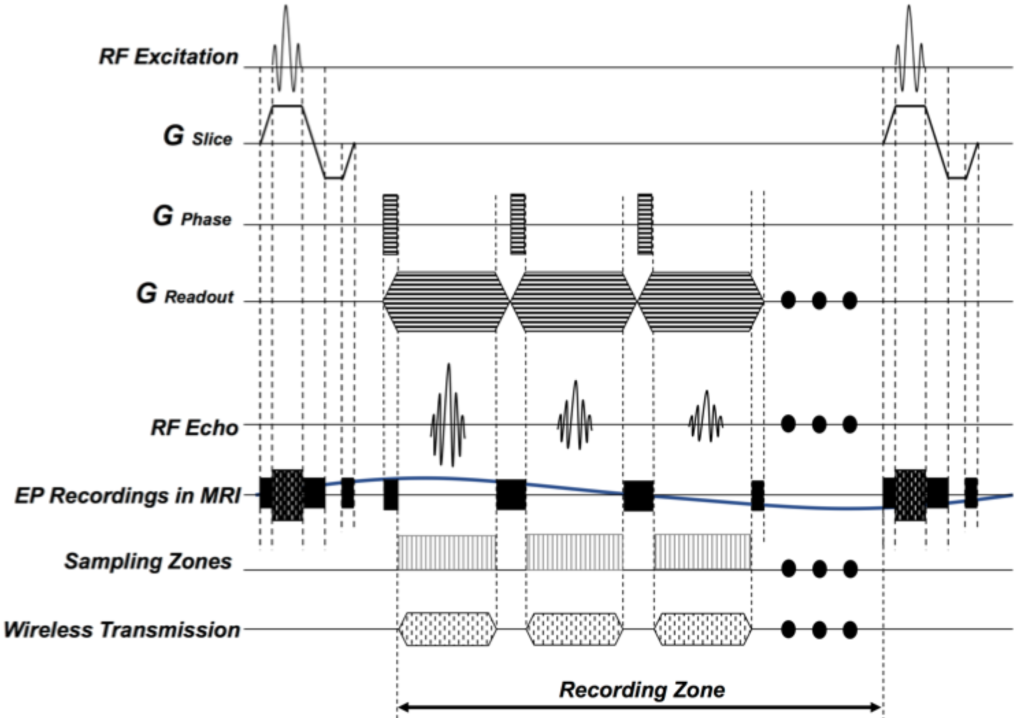
Gradient and RF pulses, as generated during the echo-planar-imaging (EPI) sequence are shown on rows 1-4. Row 5 shows the RF echo generated from the imaged subject, during the plataeu period of the readout gradient. The receiver coil of the MRI is turned on during this period. The discrete nature of the electromagnetic artifacts and there timings with respect to the gradient and RF pulses are shown on row 6. Row 7 shows the sampling zones, that avoid the EM artifacts. Row 8 depicts the wireless transmission periods, that are placed in middle of the readout zones. As a result, the transmitted data is picked up through the receiver coil along with the RF echo, emitted from the subject.

For each k-space line acquisition, the readout gradient had a plateau period of 600µs along with a ramp up and ramp down period of 100µs each. During the plateau period, 16 samples of EP signal were taken and averaged within the µC. The processed data was then transmitted in the next plateau period. 64 EP data points were recorded through the MR-image at a sampling rate of 1.3KHz for every fMRI slice acquired given the imaging sequence described in “MRI Acquisition”.

#### 6.) EP Data Extraction and Processing

Right after the scans were completed, the *fid* data (raw kspace data) was fed into a custom software, developed using MATLAB (MathWorks, MA, USA). The software isolated the MR-data and non-MR data based on their respective frequency ranges.

The scanner’s digital processing system acted as a mixer for the FSK modulated data. The isolated, non-MR data contained the wirelessly transmitted EP signal which was already demodulated at the MR-center frequency. Firstly, the “mark” and “space” frequencies (binary ‘1’ and ‘0’) were identified by calculation of power spectral density (PSD) of the non-MR data set. In the second step, asynchronous or non-coherent demodulation scheme [38], [39] was implemented to retrieve the digital packets (Figure 6). In this scheme, the received signal was treated as a sum of two amplitude shift keyed signals (ASK) and matched filters isolated these two ASK signals. Finally, the binary signal train was recovered through two envelope detectors and a logic synthesizer [39], all implemented on MATLAB.

**Fig. 6.**
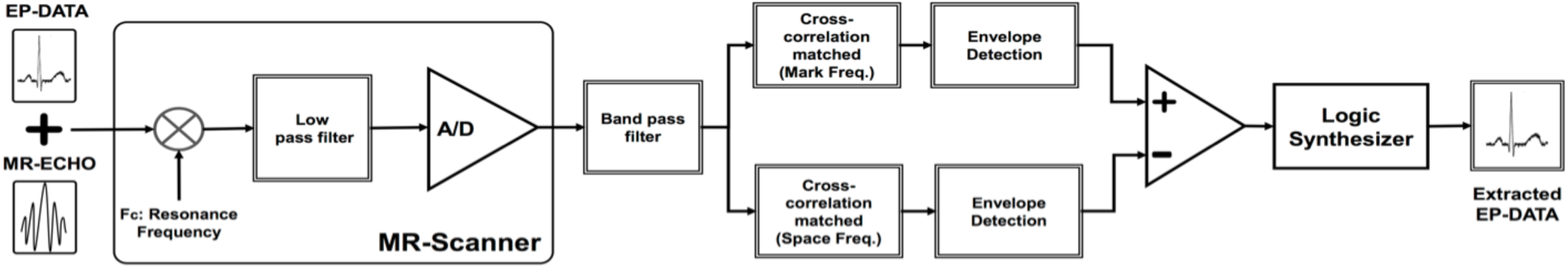
Block diagram of the wireless EP-data reconstruction method. Both MR-echo and modulated EP signal, sensed by the receiver coil is demodulated at MRI-center frequency (300.033 Mhz) and digitized by the scanner. The raw fMRI data is then bandpass filtered to isolate the non-MR data and standard FSK demodulation scheme was employed to reconstruct the device recorded EP data.

### B. Experiments

Our current investigation focused primarily on implementation of the proposed method with phantom imaging (Phantom Experiment) while simulated EP data was recorded by the device. Additionally, the efficacy of the device was evaluated through simultaneous recording of cardiac and brain electrical activity during simultaneous fMRI in rats. Finally, the effects of the device operation within MR-bore was analyzed qualitatively and quantitatively to further illustrate the advantages of the proposed technique.

#### 1.) Phantom and Animal Preparation

In the phantom study, a test tube with CuSO_4_ x 2H_2_O (1g/L) solution was used. As for the animal study, three Sprague Dawley rats (male, 350-450g, Envigo RMS, Indianapolis, IN) were used for in vivo experiments. One animal was used for LFP and SEP study; one animal was used for ECG fMRI simultaneous recording experiment; one animal was used for LFP fMRI simultaneous recording. All the experiments were performed according to a protocol approved by the Purdue Animal Care and Use Committee and the Laboratory Animal Program.

In all the experiments, animals were anesthetized following the same protocol. Each animal was initially anesthetized with 5% isoflurane for five minutes and maintained with continuous administration of 2–3% isoflurane mixed with oxygen. The heart and respiration rates and oxygen saturation level (SpO_2_) were monitored by using a small-animal physiological monitoring system (Kent Scientific, CT, USA). The animal’s body temperature was maintained at 37 ± 0.5°C using an animal-heating system.

In LFP recording experiments, electrodes and anchor screws were designed to be MR-compatible. The LFP recording electrodes were made of platinum and had a diameter of 0.127mm and a length of 3mm [40] (Plastics One, VA, USA). The reference electrode was of the same diameter with the length of 10mm. Both active and reference electrodes were made from platinum (Pt). Additionally, the pedestal of the electrodes and the inner connectors were made of plastic and copper, respectively, to be MR-compatible. To secure the electrode, nylon plastic screws were used as anchors [41]. Similar types of screws served to provide ground and reference points on the skull. A 0.3mm hole was drilled into the screws through which the ground and reference electrodes could pass to contact the brain.

Additional surgical preparation was done for LFP recording experiments. The rat was prepared for the electrode implant by administering painkiller (carprofen, 5mg/kg) before the start of the surgery. After the toe-pinch test, the rat was shaved at the site of the surgery and placed into a stereotaxic instrument (Stoelting, IL, USA). The ground and reference points were placed in the region above the cerebellum (posterior to the lambda). Nylon plastic screws [41] with holes were secured into the skull. The reference electrode and a separate ground wire were inserted into the screws. Then, a 3mm long Pt electrode [40] was implanted through a 1mm hole drilled through the skull into the right somatosensory cortex (R-S1FL) according to the rat brain anatomy [42].

For ECG recordings, the rat was fully anesthetized (3% isoflurane in oxygen at 1L/min) and shaved at the site where the electrodes were placed. An area stretching from collar of the animal down to the mid-torso was shaved to provide sufficient surface for a clean pad connection. The rat’s fur was first trimmed using a conventional trimmer, then completely removed using hair removal cream. Ag/AgCl, radiolucent ECG electrodes [43] (NeoTech Products, Inc., CA, USA) were placed in Einthoven’s Triangle formation [16], [44] for recording.

#### 2.) fMRI Acquisition

The animals were scanned in a 7-tesla horizontal-bore small animal MRI system (BioSpec 70/30; Bruker Instruments, Billerica, USA) equipped with a gradient insert (maximum gradient: 200mT/m; maximum slew rate: 640T/m/s), a volume transmit ^1^H RF coil (86 mm inner-diameter) and a surface receive ^1^H RF coil (10 mm inner-diameter). An echo planer imaging (EPI) sequence was used for fMRI scans with an echo time of 16.521ms, repetition time of 974.442ms, image size of 256×64, and a FOV of 128×32.

#### 3.) Phantom Experiment

The test tube phantom and the MR-Link device was both placed inside the MRI while the phantom was imaged using EPI. Initially, the device was supplied with a dummy signal from a function generator. The device then transmitted the amplified and digitized data wirelessly to the MR-receiver coil. The power and frequency of wireless transmitter were tuned to avoid any overlap with the imaging data.

The MR-Link transmission was once synchronized using the gradient signal from the MRI server and later utilizing the gradient detection system. This process was carried out to evaluate the efficacy of the in-built gradient detection circuit of the device. The gradient detection module was further evaluated to produce an accurate response to changing magnetic field in the MRI bore. The pick-up coil was placed at 45° angle with respect to the Gx and Gy plane to ensure magnetic variation along the x or y axis were detected by the gradient detection circuit. The coil was secured to the outside of a custom-made animal tray to keep the orientation stable throughout the experiments.

#### 4.) In vivo LFP and SEP outside MRI

Apart from recording the dummy signal, efficacy of the amplifier system was further verified through the acquisition of neural signals (LFP and SEP) from a rat, placed outside the MR-bore. LFP and SEP were recorded from R-S1FL under two conditions:
a. Variation of anesthesia levels: Isoflurane concentration was varied from below 1% (in oxygen at 0.5L/min) up to 4% to change the anesthesia level in the rat.
b. Forepaw electrical stimulation: the left forepaw was stimulated through a pair of needle electrodes. Stimulation current was delivered in 30s-ON-30s-OFF cycles through a benchtop current stimulator (Model 2200, A-M system). When it was ON, monophasic square pulses (pulse width: 5ms; amplitude: 1.0 mA; frequency: 10 Hz) were delivered. For a total of 10 mins, ten cycles of stimulation were delivered in each imaging session.

#### 5). In vivo ECG during fMRI

The in vivo ECG experiment was first conducted to monitor a reliable and well-known bio-signal to validate the device’s function inside the MRI [16], [33]. Outside of the MRI, the ECG signal was recorded by the device, as well as a benchtop amplifier (Cp511, Grass Instruments). The latter was used as a benchmark to verify that the device was functioning properly outside the MRI. Then, the rat was placed inside the MRI in supine position and the electrode leads were connected with the device. The device was placed adjacent to the rat at the foot of the animal tray holder for recording during concurrent fMRI.

#### 6.) In vivo LFP and SEP during fMRI

An experiment was conducted to monitor the evoked potential in R-S1FL of the rat during concurrent fMRI by using the same paradigm as used for SEP recording outside the MRI. Inside the MRI scanner, the rat was in a prone position. To better induce a clear hemodynamic response in R-S1FL, we used 0.03mg/Kg/h dexmedetomidine subcutaneously together with 0.2-0.3% isoflurane. The device placed was –8cm off the isocenter of the MRI for recording during concurrent fMRI.

#### 7.) Effects on fMRI data quality

The effects of wireless transmission on MR-images were quantitatively assessed through analyzing the temporal signalto-noise ratio (tSNR) of the fMRI data and the BOLD activation map with forepaw stimulation when the MR-Link device was “ON” vs. “OFF”. In the ON condition, the device recorded and transmitted the signal to the MRI receiver coil; In the OFF condition, the device was power-off but still inside the scanner.

The fMRI data were processed using Analysis of Functional Neuroimages (AFNI). The fMRI data were corrected for motion by registering every volume to the first volume using *3dvolreg*. After removing the first ten volumes, *slicetimer* was used to correct the timing for each slice. The fMRI data were then spatially smoothed with a 3-D Gaussian kernel with 0.5-mm full width at half maximum (FWHM). The fMRI time series were detrended by regressing out the linear trend voxel by voxel. To calculate tSNR for every voxel, the signal mean was divided by its standard deviation. To map the evoked activation, the stimulation blocks were convolved with a canonical hemodynamic response function (HRF), modeled as a double gamma function. The correlation between this response model and the signal at every voxel was calculated and evaluated for statistical significance (*p*<0.05, uncorrected).

## III. Results

### A. Recording synthesized EP data during phantom imaging

The phantom study outlined a repeatable experimental setup to test the device for EP recording within the MR-bore. Initially, an ECG signal (3mVpp, 1.2Hz), generated from an arbitrary function generator, was chosen as a sample EP data for this purpose. Through the MR-Link device, the ECG signal was recorded, amplified, digitized, and transmitted to the MRI receiver coil during concurrent imaging of a phantom. The wireless data appeared in the extended FOV of the image as seen by the two white lines in (Figure 7). FOV extension along the readout direction increased the image width and accommodated the non-MR data effectively without any overlap. The ECG data was then isolated and demodulated with relatively high SNR. The P wave, QRS complex, and T wave were all clearly visible in the extracted ECG signal (Figure 7).

**Fig. 7.**
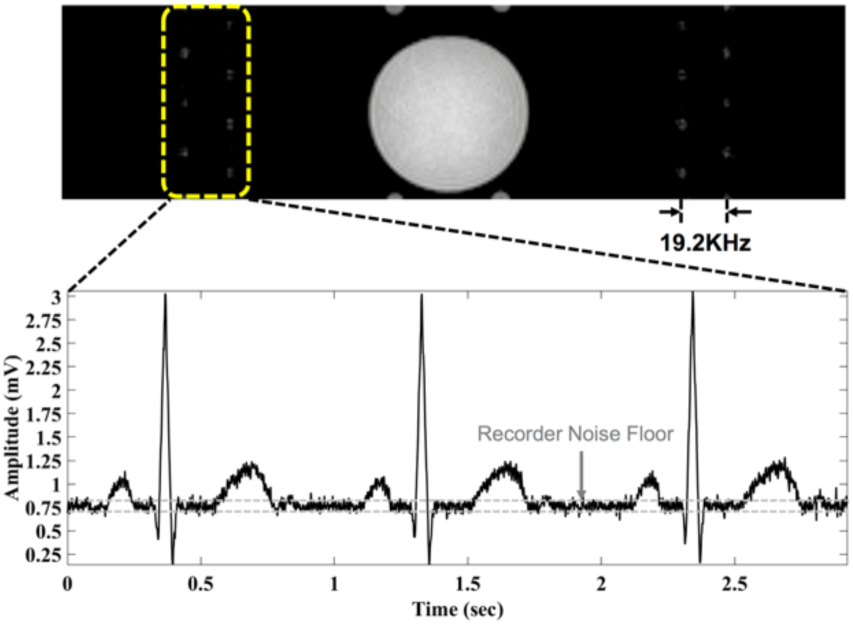
Phantom image with simulated ECG signal embedded into the FOV of MR image. The FSK modulated (BW 19.2Khz) ECG data appear as lines in the extended FOV. The plot shows the EP data extracted from the MRI.

### B. Recording LFP and SEP outside MRI

The device efficacy for in vivo recording of neural signals, e.g. LFP and SEP, was first evaluated outside of MRI. The recorded LFP varied with the level of anesthesia. As the isoflurane concentration was varied to induce deeper anesthesia, the LFP signals showed emerging characteristics of burst suppression (Figure 8). Furthermore, SEP was obtained from the signal recorded by the device by averaging as few as 10 trials (Figure 8).

**Fig. 8.**
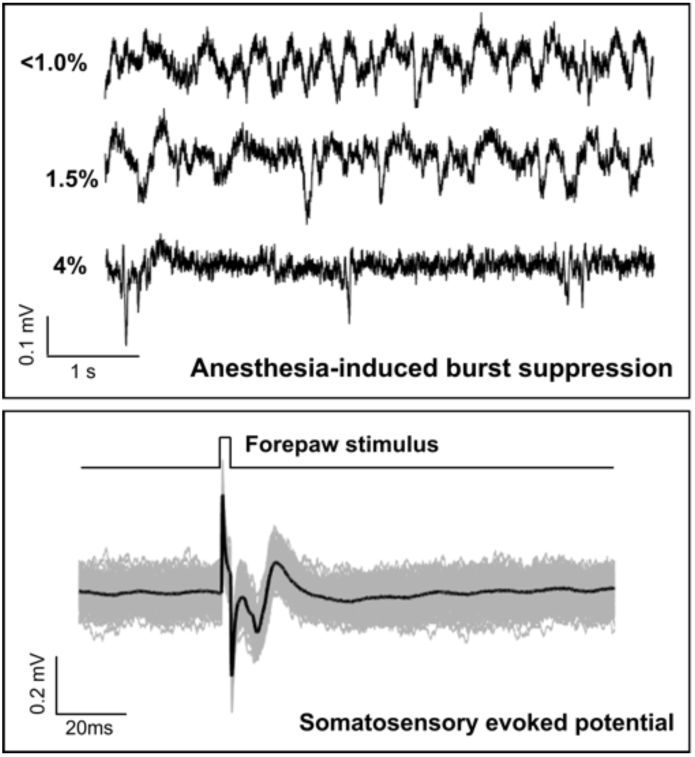
Neural signals recorded outside the MRI. Top: Spontaneous LFP varied its pattern progressively with deeper anesthesia given varying concentration of isoflurane. Bottom: SEP response to forepaw stimulation. The gray lines show the single-trial signals; The black line shows the averaged evoked response.

### C. In vivo EP recordings during simultaneous fMRI

We further tested the device for recording in vivo ECG inside the MRI scanner during concurrent fMRI acquisition. As shown in Figure 9, the device effectively rejected any gradient artifacts, while the ECG signal was extracted from the MR-images without any further post-processing. The extracted ECG data was inherently synchronized, and time stamped with corresponding brain slices. Additionally, magneto-hydrodynamic effect due to the strong static magnetic field (7T) was observed as augmented T waves [16], [45].

**Fig. 9.**
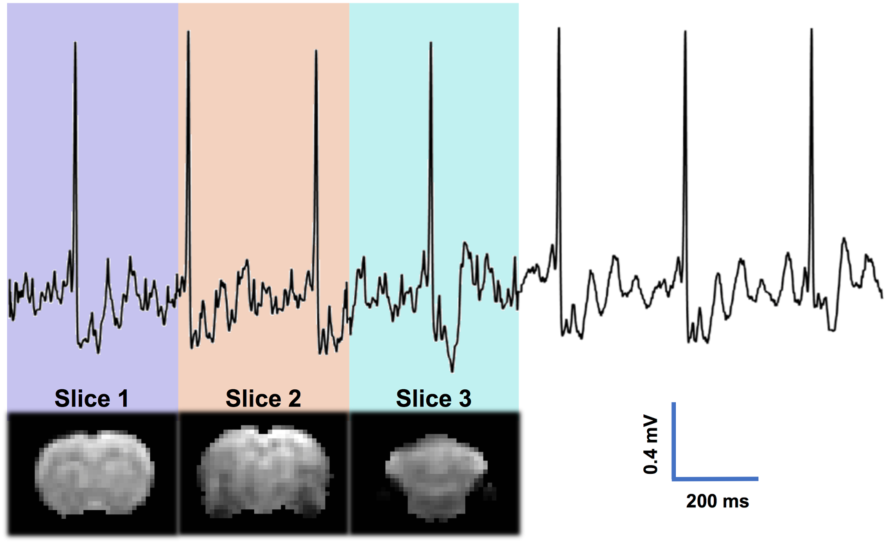
Synchronization and concurrent acquisition of ECG and fMRI.

Then we performed simultaneous fMRI and SEP recordings in vivo. As shown in Figure 10, the device successfully recorded SEP in response to the left forepaw stimulation, while the simultaneously acquired fMRI data revealed BOLD activation at R-S1FL (Figure 10).

**Fig. 10.**
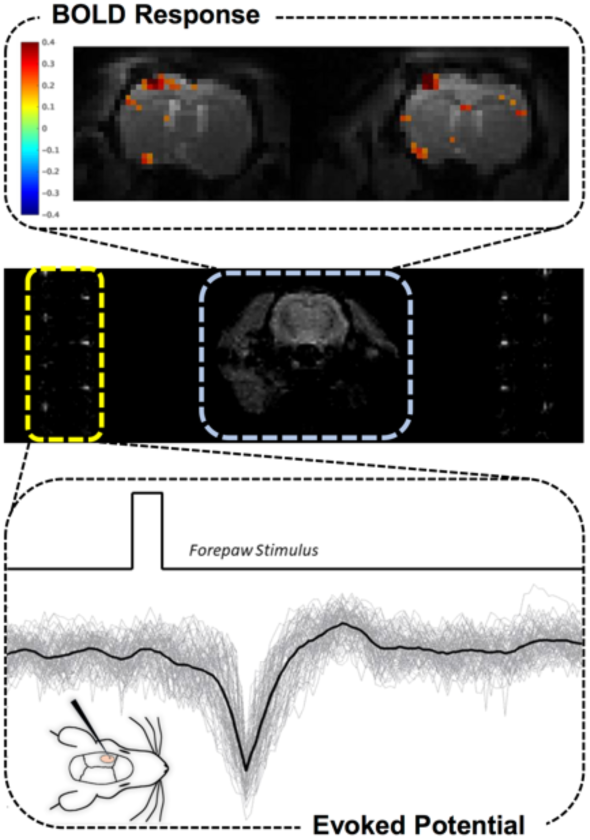
Simultaneous recording of BOLD (top) and SEP (bottom) based on the data received through MRI-receiver coil (middle).

### D. Effects on fMRI data quality

As the device transmitted EP signals in frequencies visible to the receiver coil, it was imperative to observe whether these additional non-MR signals affected the quality of BOLD signals. A systematic experimental paradigm was designed to evaluate this effect (Figure 11, top). The BOLD activation was mapped for the “ON” and “OFF” states, yielding similar results (Figure 11, right panels). Furthermore, we calculated tSNR for evert voxel. As shown in Figure 11 (left panels), the histogram of tSNR was comparable when the device was ‘ON’ vs. ‘OFF’. Thus, the device barely affected the fMRI data quality.

**Fig. 11.**
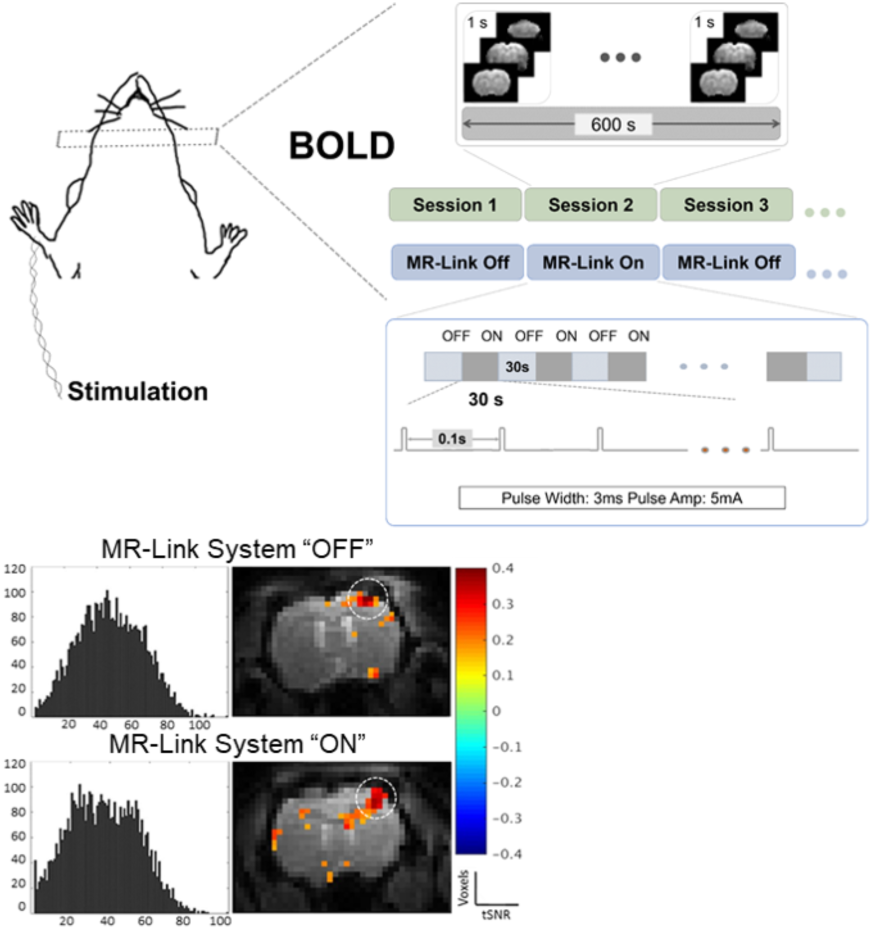
Effects of the device operation on fMRI data quality. Top: Block-design fMRI with MR-Link turned “ON” or “OFF”. Bottom: tSNR histograms (left) and activation maps (right) based the fMRI signals when the device was “ON” or “OFF”.

## IV. Discussion

The MR-Link recording system integrates with the MR-scanner and monitors its electromagnetic environment for wireless synchronization and transmission of electrophysiological recordings (Figure 2). MR-Link achieves a relatively small footprint (22 mm dia., 2gms) by utilizing the powerful digitizer and receiver capabilities of the MRI scanner. Thus, it provides an in-expensive solution for recording various types of EP signals within MRI during simultaneous imaging, without suffering from strong gradient artifacts.

### A. Utilizing MR-hardware for EP recording

The current implementation of the method only shows recording through a single channel and overlays the data on MR-images. This technique can be extended to record multi-channel EP signals. For multichannel operation, each channel’s carrier frequency can be separated into non-overlapping bands. The RF configuration in this study (receiver bandwidth: 333KHz and transmission channel bandwidth: 19.2KHz) readily allows for simultaneous recording of 9 EP channels. Additional recording channels can be incorporated by extending the receiver bandwidth and/or the FOV along the readout direction without compromising repetition time (TR).

A sampling rate of 1.3KHz, achieved during functional imaging was sufficient for slow EP signals (ECG, EEG, LFP etc.). The sampling rate was limited by wireless data transmission rate. However, application specific design of the transmitting unit and implementation of data compression techniques may increase the sampling rate, as well as the number of simultaneous recording channels significantly [46]. Time division multiplexing (TDM) or frequency division multiplexing (FDM) transmission methods can be utilized individually or in combination to accommodate higher number of channels at increased data rate.

The data extraction algorithm could work consistently as the transmission channel was placed at different distances with respect to the MR-scanner center frequency. Due to its constant carrier amplitude the digital modulation method (FSK) was more immune to noise, non-linearity and adjacent channel interference, as compared to analog modulation schemes. Moreover, the method did not require any additional signal for calibration that was needed for the implementation by Hanson et al [33]. Individual frequency channels were well separated (19.2 KHz) and detectable at very low transmission power levels (-30dBm). Furthermore, the simplicity of the processing algorithm may facilitate real-time monitoring of the EP data during simultaneous imaging through integration of the custom software with the imaging platform.

Another possible application of the method can be found with the availability of dual-tuned receiver coils. The non-MR data can be transmitted at a ‘X’-nuclear frequency without interfering with proton imaging. Our current implementation allows the MR-Link system to send the wireless data at a large range of frequencies from 75MHz to 1GHz. Additionally, the EP signals can be recorded within the MR-Link hardware and transmitted in bursts after each imaging cycle. These bursts can be picked up by MR-receiver coil through a custom designed short duration MR-pulse sequence.

### B. Power harvesting opportunities

Powering the recording system within the MR-environment requires significant attention since additional powering circuitry may potentially affect the operation of the MR-scanner. The low-powered, battery operated recording and transmission system, presented in the study may be well suited to be powered through a variety of different approaches. One such attractive method is wirelessly powering the device by harvesting energy from the strong magnetic flux generated by MRI [47]. Application of wireless powering can remove additional cables running through the MRI bore and thus may improve the safety and functionality of the device within MRI. The advantages of wireless powering provide a strong case for its implementation into the MR-Link device to further make it into a stand-alone and user-friendly system.

### C. Future development and application

The MR-Link device works with various fMRI paradigms, while requiring minimal modification to the MRI hardware and software environment. However, currently the device can only function when it is used with EPI sequences administering trapezoidal gradient waveform. The current strategy for adaptive and discrete sampling, amplification, and transmission relies on the assumption that gradients are absent or constant during certain periods in the pulse sequence. This assumption, although valid for trapezoidal gradient waveforms, is not satisfied for some pulse sequences, e.g. for spiral imaging. Future development is needed to extend the device’s operational strategy for usage with other types of pulse sequence.

The efficacy of the MR-Link recorder was tested using a 7T animal MR-scanner with gradient amplitude and maximum slew rate of 200mT/m and 640T/m/s, respectively. These values are considerably larger than the clinically used 1.5T or 3.0T MRI with gradient strengths up to 45mT/m and a slew rate up to 200mT/m/s. As a result, the gradient artifacts generated with our test setup were considerably larger than that, anticipated for a human MRI system. Furthermore, given MR-Link’s capability to avoid EM artifacts, the extracted data did not require any additional retrospective digital processing [11], [20], [48]. The small footprint of this device also renders itself safer for placement inside MRI. Taken together, the method presented herein is expected to be readily usable with human MRI. Therefore, the success of the current research is expected to open new avenues for widely accessible and integrative neuroimaging tools.

## Notes

This work was supported in part by NIH R01MH104402, R41NS105298, Purdue Research Foundation and Purdue University. Authors have no conflict of interest.

